# Networks of marine protected areas under fluctuating connectivity: The importance of within-MPA dynamics

**DOI:** 10.1101/2020.12.05.413013

**Authors:** Ridouan Bani, Tianna Peller, Justin Marleau, Marie-Josée Fortin, Frédéric Guichard

## Abstract

The design of marine protected areas (MPAs) has been optimized under assumptions of spatially and temporally homogeneous larval dispersal, despite complex spatiotemporal patterns displayed by ocean currents. Here we studied the effect of dispersal variability on the effectiveness of MPA networks across scales. We adopted a nested approach integrating the dynamics of both within and among MPA connectivity into a stochastic metapopulation model and first derived metapopulation persistence (required reproductive effort) and stability over MPA networks by partitioning within and among MPA contributions in relation to the spatial resolution of within-MPA connectivity. We applied this framework over a range of dispersal traits (spawning time and pelagic larval duration) and MPA network configurations, based on simulated biophysical connectivity along the northeast Pacific coast. Our results show how within-MPA dynamics affect predictions based on parameters of MPA networks such as MPA size, spacing, and pelagic larval duration. Increasing within-MPA spatial resolution predicted increasing population persistence and stability independently of other network properties. High-resolution within-MPA dynamics also predicted a negative relationship between species persistence and MPA spacing while that relationship was non-monotonic under low-resolution within-MPA dynamics. Our analysis also resolved the role of pelagic larval duration for scaling up within-MPA dynamics to MPA networks: species with short larval duration led to increasing network stability with MPA spacing while the opposite was observed for species with long larval duration. Our study stresses the importance of integrating fluctuating larval connectivity, both within and among MPAs, and more specifically suggest the benefit of small and nearby MPAs under increasing ocean variability.

## Introduction

Networks of marine protected area (MPA) are designed to allow for synergistic interactions among individual MPAs across spatial scales and for the protection of biological diversity, and of natural and associated cultural resources (IUCN 1994; WCPA/IUCN 2007). Effective MPAs network planning can allow populations to grow and persist within MPAs and contribute larval recruitment (Harrison et al. 2012; Baetscher et al. 2019) as well as spillover through adult movement (Kellner et al. 2008; Moffitt et al. 2009; Goñi et al. 2010; Kough et al. 2019) to unprotected areas. Metapopulation theories concerning MPA networks associate these principles with four proxies: location (e.g. representation, nurseries, special features), size (e.g. richness, adult movement, home range), spacing (e.g. connectivity), and/or number (e.g. replication). Effective MPA network planning is equivalent to understanding how location, size, spacing and number of MPAs contribute to species reproductive (Goodyear 1993) and dispersal (Moffitt et al. 2011; Green et al. 2015) abilities. Many studies concerned with principles for designing MPAs networks that address social, economic and biological criteria, but few have set these principles in the context of variability in environmental conditions and biological traits (McLeod et al. 2009; Magris et al. 2014). MPA network design needs to reflect the complexity of species’ niches, especially the spatial configuration and environmental and biological spatiotemporal variability (Caswell et al. 1997; Caswell 2001; Lande 2002; Tuljapurkar and Caswell 2012; Tuljapurkar 2013).

Theoretical studies concerned with the effects of MPA size and spacing on species persistence show that MPA size and the fraction of coastline protection required for species persistence should be proportional to mean dispersal distance (Botsford et al. 2019). However, others have drawn attention to the impact dispersal heterogeneity can have on species persistence in MPAs (Kaplan 2006; Aiken et al. 2007; Kaplan et al. 2009; White et al. 2010; Aiken and Navarrete 2011). Recent studies have indicated the importance of larval dispersal patterns, especially the spatiotemporal variability that can arise from hydrodynamic processes, climatic variability, and from developmental and behavioral changes (Munday et al. 2008; Siegel et al. 2008; Mitarai et al. 2009; Munday et al. 2009; Lett et al. 2010; Magris et al. 2014). Current MPA network theories typically ignore such variability by describing larval transport and development with dispersal kernels (Gaylord et al. 2006; Aiken et al. 2007; Woodson et al. 2012; Kerr et al. 2015; Le Corre et al. 2019). Spatiotemporal fluctuations and aggregation in recruitment events can be observed over a broad range of spatial scales, including scales that are smaller than the size of an MPA. This can lead to a network of larval connectivity within individual MPAs in addition to among-MPA connectivity.

Spatiotemporal variability in dispersal can produce inflating or deflating effects on growth and density (Watson et al. 2012; Williams and Hastings 2013) which may affect stability of metapopulation in MPA networks (Bani et al. 2019) and can be used in MPA network planning especially in extreme conditions when average larval recruitment is low. These inflating or deflating effects are purely a product of dispersal fluctuations and depend on the temporal pattern of larval exchange among populations, a feature that dispersal kernels fail to address when modeling dispersal dynamics. When MPA network connectivity is assumed static and larval recruitment fails to offset post-settlement mortality, the metapopulation defined by the MPA network will go extinct. Positive growth and stable dynamics can still be achieved when fluctuations in larval recruitment are autocorrelated (Williams and Hastings 2013) and spatially asynchronous (Watson et al. 2012; Bani et al. 2019). Recent theory applied to simulated biophysical connectivity (Bani et al. 2019), predicts the species density in MPA networks can undergo these inflating or deflating effects with changes in the configuration of MPA networks (spatial heterogeneity in larval recruitment, spacing, proportion of protected area). However, there is currently no theory of how these effects of spatiotemporal patterns of connectivity can scale up from within population dynamics to whole metapopulations. Such theory is needed to inform the optimal configuration of MPA networks under increasing climate and ocean variability.

Here, we adopt a metapopulation model (Bani et al. 2019) that captures the dynamics of a single marine species subject to density regulation and occupying MPAs interconnected by time varying larval dispersal. We predict the growth and stability of the metapopulation in a MPA network from the stochastic and stage-structured estimates of stationary distribution. We derive an explicit expression for metapopulation stability as a contribution of scale-dependent effect of within- and among-MPA fluctuations in larval recruitment. We then apply our framework to the Pacific northeast coastal system, where we vary the spacing, and proportion of protected area by either random or structured MPA assignment along the coast.

## 1 Metapopulation dynamics within MPA networks

We study populations dynamics of a single marine species characterized by an adult sedentary phase and a dispersive larval phase. The populations are distributed over an MPAs network noted 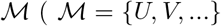 is the network of MPAs, *U*, and *V* are two individual MPA within the network 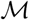, and 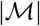 is the number of MPAs). The population occupying the MPAs ({*U, V*,…}) are interconnected by time-varying larval dispersal. We adopted a stochastic and density-dependent model that assumes densities are near carrying capacity. Our stochastic framework captures and predict species metapopulation growth and stability conditions when dispersal varies in time and space.

The population *N_U_*(*t*) within each MPA at time changes due to three processes: (1) adult individuals die with a per-capita rate *m_a_*, (2) a time constant self-recruitment with a per-capita rate of 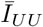and (3) time varying externalrecruitment from other MPAs 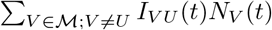. We used a time averaged value for local recruitment (local-retention) because larvae that recruit directly back to their natal population are not exposed to ocean current variability compared to larvae that travel greater distances. We also assume that the intensity of each process varied with local population size to account for competition among conspecifics at both recruitment and adult stages. Competition can be for space or for other resources, and is implemented as a logistic function with carrying-capacity *K_U_* specific to each MPA.

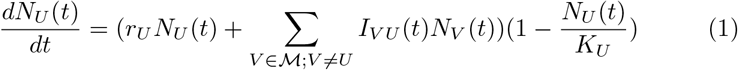

The local growth is equal to the adult mortality rate subtracted from the self-recruitment rate 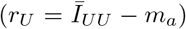. The time varying per capita immigration rate *I_VU_*(*t*) = *δfC_VU_*(*t*) is a function of *C_VU_*(*t*): the probability or proportion of individuals dispersing from *V* to *U* at time *t*, *f* and *δ* are adult fecundity and larval survival rates respectively and are spatially uniform over the metapopulation. It is important to note that the loss of larvae to unprotected areas may be of importance for fisheries purposes, however, our model only tracks the dynamics of population’ individuals within MPAs and we assume no survival of larvae (*δ* = 0) outside MPAs.

We convert the density-dependent model in eq. (1) to a stochastic and multivariate Markovian process that converges to a stationary distribution when its deterministic counterpart reaches the stable equilibrium (*K_U_, K_V_*,…). If we assume that all the MPAs have similar relative carrying capacity (i.e. 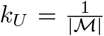), the mean and variance of the stationary distribution of regional relative abundance are as follows (Bani et al. 2019):

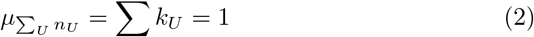

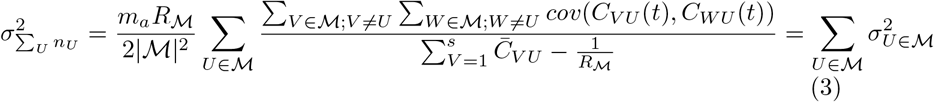

### 1.1 Metapopulation stability

We can consider the variation in total relative abundance over the MPA network around carrying capacity as a measure of population stability defined as:

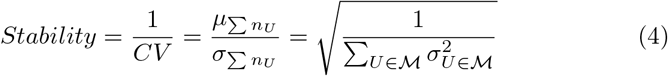

The contribution of each MPA to MPAs network’ metapopulation stability measured in terms of the variance contribution of each MPA to metapopulation variance, which is represented by the variance of the stationary distribution 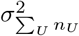 (eq. 3). For instance, the contribution of MPA U is:

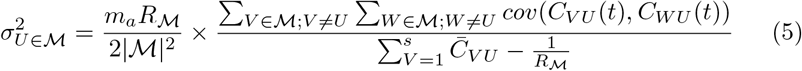

The variance contribution (eq. 5) assumes spatially homogeneous larval recruitment within MPAs. We now develop an expression for the variance contribution of each MPA to MPAs network? stability for the case where within-MPA larval recruitment can be heterogeneous.

### 1.2 Minimal net reproductive rate for persistence

For the populations occupying the MPAs network and following the dynamics described in eq. (1), it is sufficient to start with a population size close to the carrying capacity for the dynamics to converge toward the stationary distribution (eqs. 2 and 3). However, in reality, there may be disturbance events, such as extreme environmental conditions that cause sudden drops of abundance. To account for such events, we imposed an additional condition to allow the species abundance to grow back and remain close to the carrying capacity.

At low abundance (a.k.a. 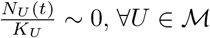), we optimized the fecundity (*f_opt_*) such that the largest eigenvalue (λ_*d*_) of the average projection matrix of the density-independent (a.k.a. average connectivity matrix - diagonal adult mortality matrix) version of model (eq. 1) to be equal to 1, and we define the minimal net reproductive rate (NRR) of the MPAs network as 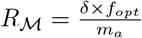.

Our minimal net reproductive rate 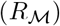 required for persistence differs from the net reproductive rate *R*_0_ defined as the average number of recruits into the first age class produced during an individual’s life (Caswell 2001).

This is because, (*i*) we use stage-structured classes representing populations rather than age classes, (*ii*) incorporates the optimized connectivity matrix to balance adult mortality, and (*iii*) metapopulation in MPAs network can persist if net reproductive rate is higher than its minimal value required for persistence 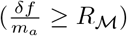 while *R*_0_ > 1 is required for persistence.

For the single population/MPA case, the minimal net reproductive rate is equal to the inverse of local-retention 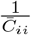, where 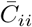 is the time-averaged proportion of larvae recruiting back to natal population.

For a metapopulation to persist in a MPAs network, the net reproductive rate should be larger than the minimum value required for the densities to grow back and converge toward the stationary distribution with a stable fluctuation (i.e. smaller variance). We now study the dynamics of a metapopulation in MPA networks when larval recruitment is spatially heterogeneous within individual MPAs.

## 2 Spatial resolution of larval recruitment within MPAs

In addition to the temporal variability in immigration terms (*I_VU_*(*t*)), we also consider the case of spatially heterogeneous larval recruitment within individual MPAs. In such case, we divide each MPA U into smaller patches (*P_i_*) such that *U* = {*P*_1_, …,*P_u_*}, and that within each patch *P_i_* larval recruitment is assumed to be homogeneous. However, we assume that larval recruitment can differ among the patches within-MPA.

The connectivity between MPAs represent a spatially averaged connectivity between within MPAs patches *P_i_*. For example, the connectivity from MPA *V* = {*P*_1_,…, *P_v_*} to *U* = {*P*_1_,…, *P_u_*} at time *t* is:

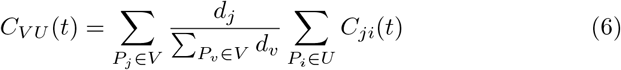

The connectivity *C_ji_*(*t*) is the proportion of the total number of larvae *d_j_* released from patch *P_j_* ∈ *V* arriving to patch *P_i_* ∈ *U*.

The size of patches *P_i_* corresponds to the level of larval recruitment spatial resolution within each MPA *U*: when MPA *U* has a spatially homogeneous larval recruitment then the size of patches *P_i_* within which larval recruitment is assumed homogeneous should be larger than the MPA *U* size (i.e. there exists a patch *P_u_* such that *U* ⊂ *P_u_*), otherwise the size of patch *P_i_* should be smaller than the MPA *U* size for the MPA *U* to have spatially heterogeneous larval recruitment (i.e. 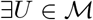 such that 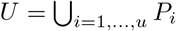 and *u* > 1).

When patches *P_i_* are smaller than the size of individual MPAs, larval connectivity can become nested within individual MPAs. The within-MPA spatial resolution in larval recruitment can affect how individual MPAs contribute to metapopulation growth and stability. In the following section, we derive an explicit relationship that partitions the contribution of MPA *U* to metapopulation stability in relation to the spatial resolution of larval recruitment within MPA *U*.

### 2.1 Partitioning MPA and within-MPA contributions to population stability

When larval recruitment is spatially heterogeneous such that patches *P_i_* are smaller than the size of individual MPAs (i.e. 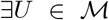 such that 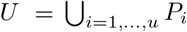 and *u* > 1), individual MPAs become themselves networks of their within patches *P_i_*. Thus, individual MPAs are subnetworks within the MPA network and their variance contribution to the MPA network metapopulation stability depends on within-MPA dynamics. We can derive the nested MPA contribution to metapopulation stability across the whole MPA network starting from MPA *U* contribution expression with spatially homogeneous larval recruitment (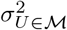 in eq. 5) and by integrating between-MPA connectivity as a function of within-MPA patches *P_i_* connectivity (eq. 6) (see appendix B for details).

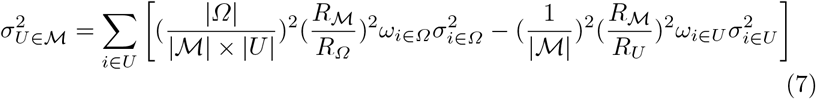

The population within patch *P_i_* contributes

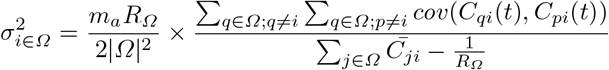

to whole 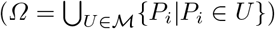 metapopulation variance, and

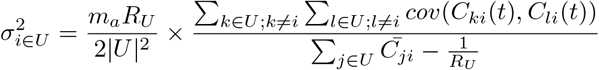

to within MPA *U* ({*P_i_*|*P_i_* ∈ *U*}) metapopulations variance (Fig. 1). Both contributions are weighted by

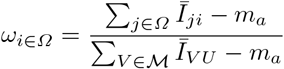

and

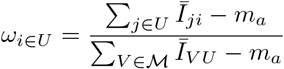

which are patch *P_i_* larval recruitment relative to adult mortality when recruitment is from the whole Ω or from within-MPA *U* metapopulations, respectively. 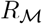 *R_Ω_*, and *R_U_* are the net reproductive rates of MPAs network 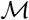 *R_Ω_* when larval recruitment is spatially homogeneous within MPAs, of whole metapopulation *Ω* across patches *P_i_* (regardless of the MPA they belong to), and of patches within-MPA *U*.

**Fig. 1.**
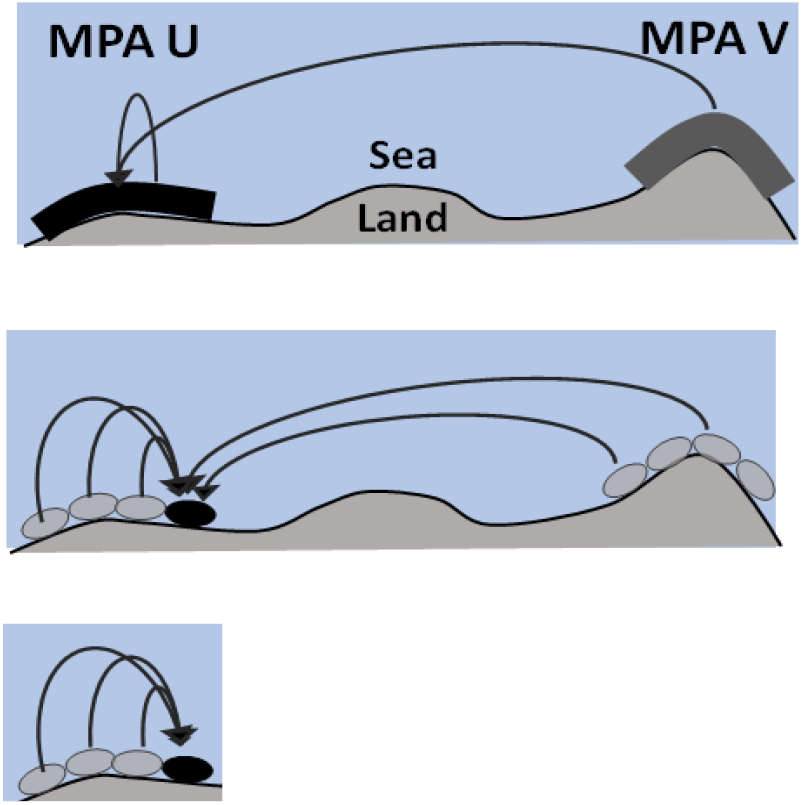
An illustration of two MPAs (top panel: 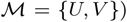) with equal size when larval recruitment is well mixed within both MPAs and showing two connectivity pathways: a self-recruitment back to MPA *U* and external recruitment from MPA *V*. When larval recruitment is spatially heterogeneous, MPAs become sub-networks (middle panel) and the both connectivity arrows shown for well-mixed MPAs (top panel) are replaced with multiple connectivity arrows between patches. Spatial heterogeneity in larval recruitment can generate within-MPA dynamics (bottom panel).

It is important to note that while 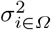 measures variance contribution of each *P_i_* population to stability of metapopulation within all MPAs (*Ω*), 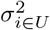 measures the contribution of the same population to stability of metapopulation within-MPA (*U*) only (Fig. 1). The right-hand side of eq. (7) is composed of two parts, the first part measures the weighted contribution of each patch to the metapopulation of all (*Ω*) regardless of what MPA they belong two and the second part measures its contribution to the metapopulation within the MPA it belongs to.

We verify that the relationship in eq. (7) holds for two boundary conditions when larval recruitment is homogenous within-MPA (i.e. 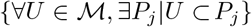) as well as for the case of a single MPA. When larval recruitment is homogenous within-MPA, 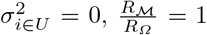, and *ω*_*i*∈*Ω*_ = 1 and MPA variance contribution in eq. (7) reduces to the case when larval recruitment is homogenous within-MPA in eq. (5). Moreover, in the case of a single MPA, 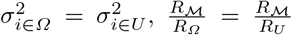, and 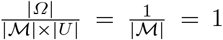 such that 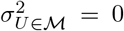 regardless of the level of within-MPA spatial resolution. In this case, single MPA dynamics with homogeneous larval recruitment are governed by local growth rate 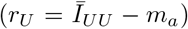 and since self-recruitment is optimized to balance adult mortality, the population grows deterministically to reach carrying (*i.e*. 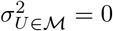).

By using the nested network contribution of MPAs to variance 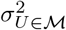 given by eq. (7) in our measure of stability (eq. 4), the stability of a network of nested MPAs becomes:

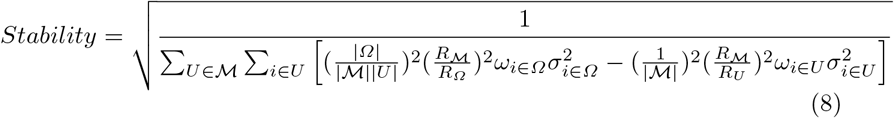

## 3 Case study along northeast Pacific coasts

### 3.1 biophysical connectivity

We apply our model to a case study along the coast of British Columbia, Canada and Washington State, USA. To quantify the dynamic connectivity, a solution from a three-dimensional hydrodynamic model ROMS (Regional Ocean Modeling System) output is used to generate neutrally buoyant particle trajectories along the coast. The ROMS model uses 3 *km* horizontal resolution and a 31 vertical resolution levels. To hind-cast ocean currents over the 10-year period from 1998 to 2007, Masson and Fine (2012) forced the ROMS output with tides, 3-hourly winds from the North American Regional Reanalysis (NARR; Mesinger et al. (2006)), heat fluxes, freshwater runoff (Morrison et al. 2012), and lateral boundary salinities and temperature from Simple Ocean Data Assimilation (SODA; Carton and Giese (2008)).

444 circular release sites (5 *km* radius) with centers located on the coastline are distributed uniformly throughout the nearshore (Fig. 3-a). 133 million passive Lagrangian particles (LTRANS: North et al. (2011)) were released from the 444 patches for a period of 10 years between *Jan* 1, 1998 and *Aug* 26, 2007. The passive particles were released biweekly starting *Jan* 1 to *Aug* 26 every year and then tracked (Latitude, Longitude) weekly for a period of 18 weeks (120 days). We refer to the release day by spawning time and the duration spent after release as pelagic larval duration, so we can assume hypothetical species represented by pairs of spawning time and pelagic larval duration. In total we had 18 spawning times ({Jan 01, Jun 15,…, *Aug* 12, *Aug* 26}) and 18 pelagic larval durations ({1, 8,…, 113,120}) and a total of 18 × 18 = 324 hypothetical species. We assume that our hypothetical species reproduce during the one-day spawning time window every year and that all larvae reach competency to settle exactly after the pelagic larval duration. Connectivity among the 444 circular patches are probability of moving from a patch to another and are calculated as the proportion of particles that are within the 5 km radius of each patches divided by the number of particles released from the source patch. The connectivity times series are calculated for each pair of spawning time, pelagic larval duration and year.

### 3.2 MPA network configuration: MPA size, MPA spacing and within-MPA connectivity

We studied the effects of within-MPA connectivity and of its spatial resolution on MPA persistence defined as the minimal net reproductive rate (minimum NRR) and stability 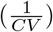. We assumed that and individual MPA can be formed by aggregating neighboring sites of 5 km radius (Fig. 2a). We first assessed the effect of spatial resolution on local retention and minimal net reproductive rate for the case of single MPAs by aggregating contiguous sites into a single patch forming a single MPA of increasing size, up to 100% of the total 444 sites. MPAs of each size were replicated by varying the location of the starting site along the coastline.

**Fig. 2.**
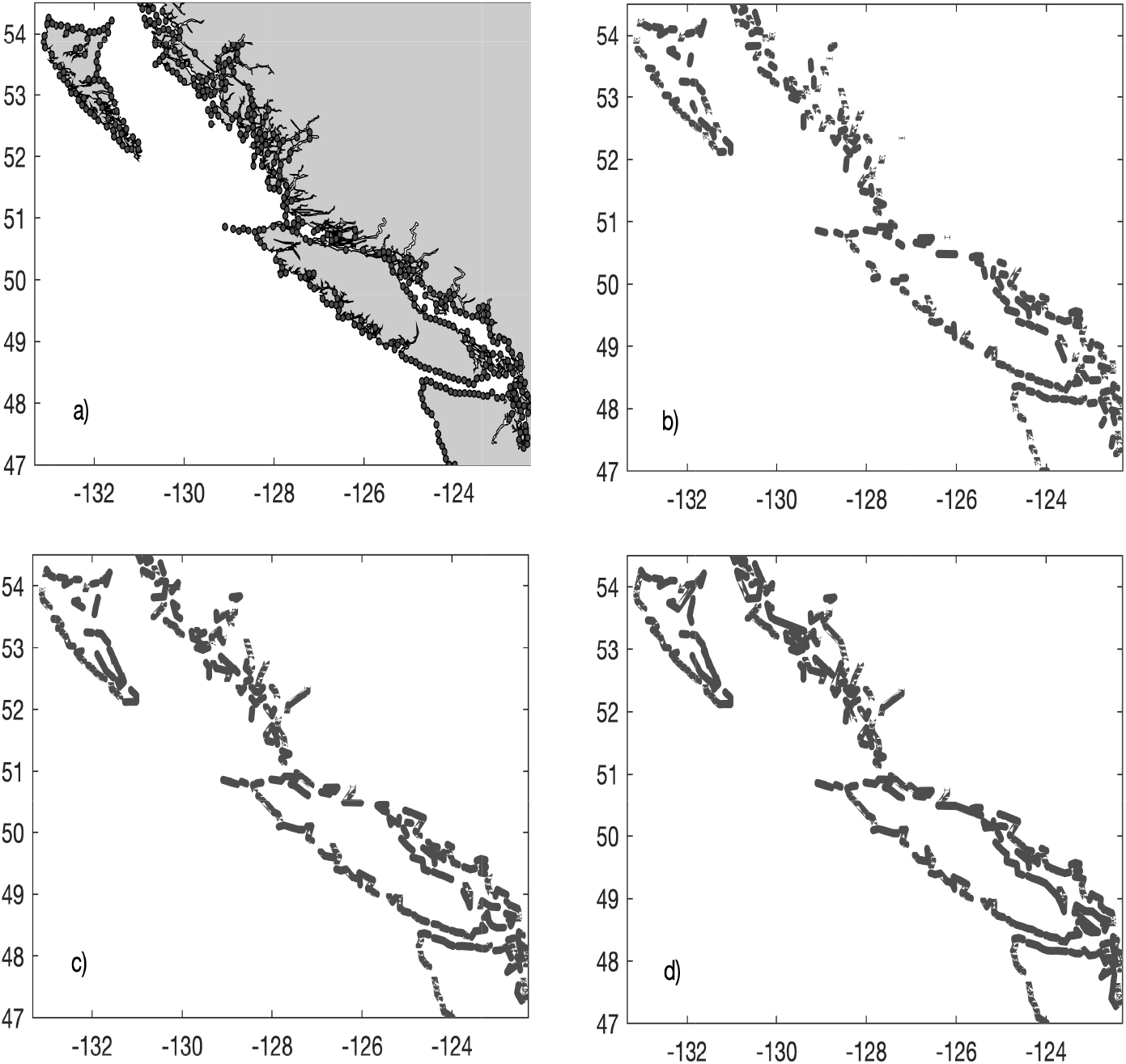
The North-East Pacific coast (British Columbia, Canada and Washington, U.S.) and the distribution of (a) 10 km sites along the coastline, and aggregated patches of (b) 2-times, (c) 4-times, and (d) 8-times the size of circular patches. The patches: 1-time, 2-times, 4-times, and 8-times the size of the site with 10 km diameter are the four levels of spatial heterogeneity in larval recruitment within-MPA. We used equation (6) to calculate the connectivity between the patches using the connectivity between the sites.

We implemented MPA networks and controlled for MPA spacing by limiting the size of individual MPAs to an area equivalent to 8 contiguous sites (80 km), with a total of 49 MPAs (Fig. 2d). We varied the proportion of protected area (10%, 20%, 30%, and 40%) by randomly sampling (5, 10, 15, and 20) out of 49 potential MPAs. We further divided the sampled MPAs into 5 categories of MPA spacing (76 km, 163 km, 250 km, 337 km, and 425 km), resulting in the same number of MPA network replicates for all distance categories. The MPA spacing was calculated as an average distance between MPAs as follows 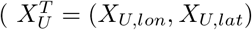 is the center of MPA *U*):

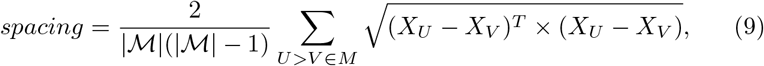

For example, if we assume a network of 5 MPAs, the MPA spacing is calculated as follows:

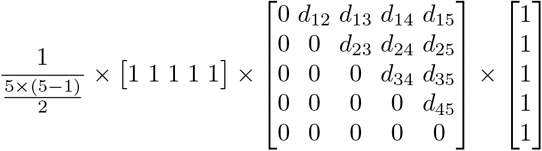

Where 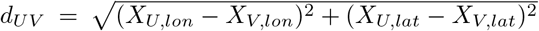 is the distance between MPA *U* and *V*. When applied to the case study, the MPA spacing range span between 32 km and 469 km and 76 km, 163 km, 250 km, 337 km, and 425 km are representative values of the 5 different distance groups.

To control for within-MPA spatial resolution (*i.e*. size of patches *P_i_*), we aggregated sites within-MPAs from its highest level of spatial resolution corresponding to a single site (no aggregation) to patches of increasing size corresponding to 3 lower spatial resolution levels represented by patch length: 10 km (1-site in Fig. 2a), 20 km (2-site aggregation in Fig. 2b), 40 km (4-site aggregation in Fig. 2c), and 80 km (8-site aggregation in Fig. 2d). Within-MPA, larval recruitment is spatially-homogeneous when the spatial resolution is as coarse as MPA size. Within-MPA spatial resolution increases with decreasing patch length.

## 4 Results

### 4.1 Local retention and the role of MPA size and larval duration

Under the assumption of homogeneous larval dispersal over MPAs, local retention defined as the proportion of larvae that recruit back to natal patch (local retention) should be spatially-homogeneous per unit area, and thus independent of the location and size of the patch *P_i_* (Fig. 2). However, our results revealed that local retention is heterogeneous across patches even with only 0.2% of areas protected and that it increased with the proportion of protected areas (Fig. 3c). Average (standard deviation) local retention when sites are aggregated into patches (*P_i_*) of length 10 km, 20 km, 40 km, and 80 km is 35% (51.41%), 7.27% (7.71%), 13.37% (16.36%) and 18.84% (17.76%), respectively. Increasing the proportion of protected areas by aggregating neighboring sites/patches to form larger single MPAs (Fig. 3a) showed that local-retention followed a non-monotonic response to proportion of protected area (Fig. 3c). The minimal net reproductive rate (NRR), which is equal to the inverse of local-retention, slightly increased with the size of single MPA before it starts to decrease at 3.6% (area stretching 160 km along the coast) and even decreased faster after 14.4% of the total area (Fig. 3b: Single MPA).

**Fig. 3.**
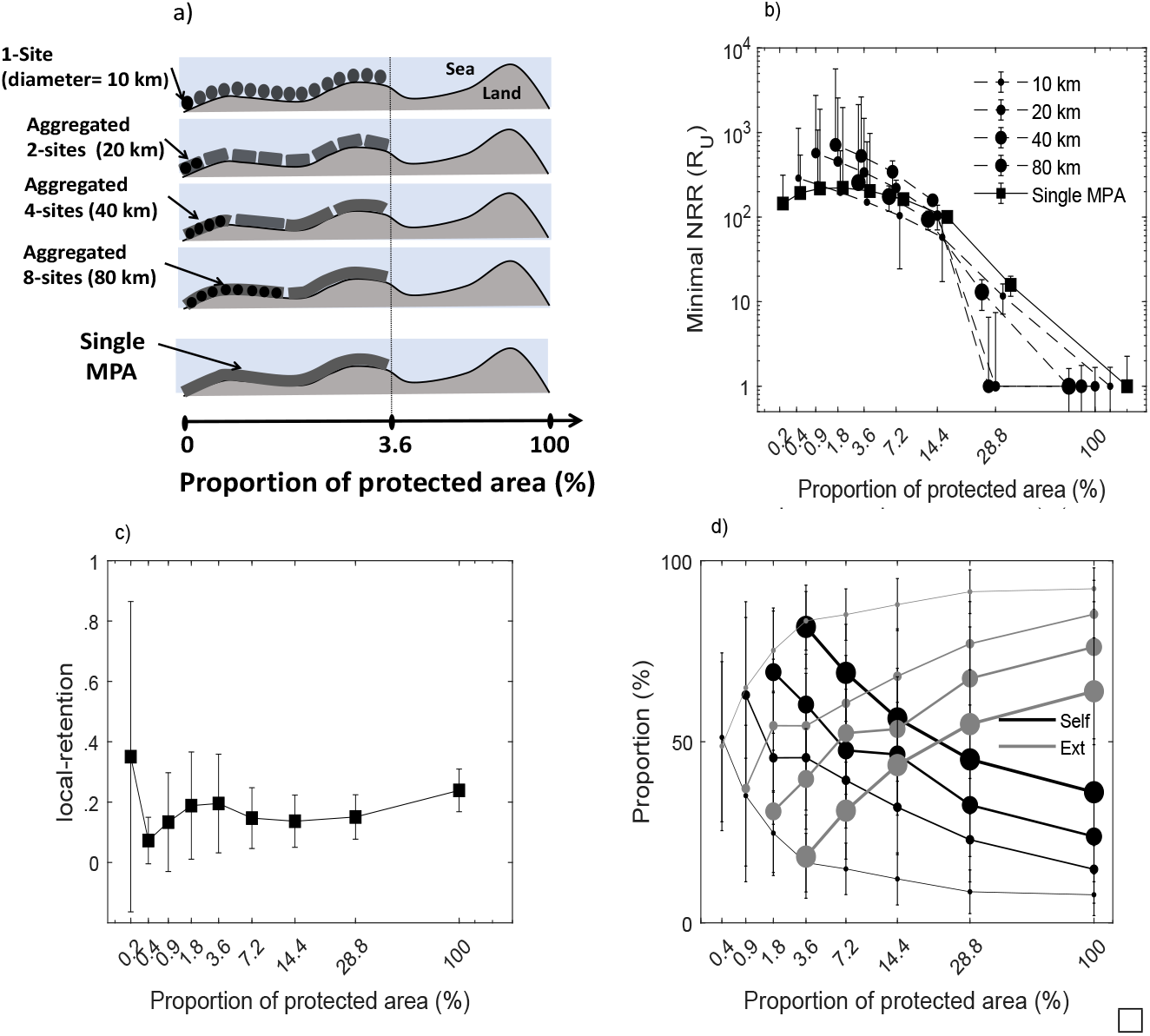
(a) Illustration of how protected 10 km sites (top panel) are aggregated into patches defining the spatial resolution of within-MPA connectivity: the resolution is highest when larval recruitment is well-mixed within 10 km sites (top), and decreases when averaged over patches of 2 (20 km), 4 (40 km) and 8 (80 km) aggregated sites. (b) Effect of the proportion of protected sites on the Minimal net reproductive rate (NRR) across values of within-MPA spatial resolution, (c) the probability of local-retention with well-mixed within-MPA larval recruitment, and (d) the proportion of self- (*Self*) and external-recruitment (*Ext*) among patches within-MPA across values of within-MPA spatial resolution. Points and error bars represent the mean and standard deviation.

Spatial variability in within-MPA local retention is stronger for species with short compared to long pelagic larval duration (Fig. B1 in Appendix B). Local retention varied little (12%) for pelagic larval duration of 120 days compared to 63.34% to 22.12% of variations when the proportion of protected area increased from 0.2% to 7.2% for pelagic larval duration of 8 days. Although spawning time has overall little effect on local-retention (Fig. B1), its effect can be comparable to the effect of pelagic larval duration under low (≤ 0.4%) proportion of protected area (Fig. B1), it can be important as pelagic larval duration for intermediate and larger proportion of protected area (≥ 7. 2% in Fig. B1). The timing of spawning can be more important than pelagic larval duration for very small MPAs while the opposite was observed for larger MPAs.

### 4.2 Within-MPA connectivity fluctuations can promote metapopulation persistence in MPA networks

Patches within MPAs form subnetworks within MPA networks. The within-MPA spatial resolution, or scale over which larval recruitment is averaged (size of patch *P_i_*) can strongly affect the predicted minimal net reproductive effort (NRR) for persistence. Our results revealed that when the proportion of protected area is higher than 14.4%, the minimal NRR of an individual MPA with spatially-homogeneous larval recruitment is larger than the minimal NRR of their nested subnetworks regardless of their level of spatial resolution (10 km, 20 km, 40 km, or 80 km; Fig. 3b). Moreover, the minimal NRR required for persistence decreased with increasing spatial resolution when the proportion of protected area is under 14.4% (Fig. 3b).

Single MPAs have more chances to persist with spatially heterogeneous compared to homogeneous larval recruitment when local retention is not sufficient to compensate for adult mortality. Self-persistent individual MPAs require a local retention high enough to compensate for adult mortality and persistence of within-MPA subnetworks requires two conditions: (1) at least a few self-persistent patches (2) that are well connected to other patches within the subnetwork. Larval recruitment spatial variability indicated that there exist many smaller 10 km (few large) patches within-MPA with high (low) local-retention that is higher than patches of 20 km, 40 km, and 80 km, and this number decreased with the size of patches (Fig. 3d), and species pelagic larval duration (Fig. B1). Thus, for patches within-MPA, the chances of self-persistence increased with the decrease of their size. Moreover, when we compared the levels of larval contribution within 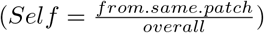 and between 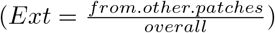 subnetworks, our results revealed that the proportion of protected area increases and decreases the proportion of external- and self-recruitment, respectively, while within-MPA spatial resolution decreases and increases the proportion of external- and self-recruitment, respectively (Fig. 3d).

External recruitment surpassed self-recruitment when within-MPA spatial resolution is high (10 km; Fig. 3d), however, the balance switched to self-recruitment as spatial resolution decreased (80km; Fig. 3d). The proportion of protected area at which the self- and external recruitment have similar contribution decreased with spatial resolution, especially for species with short pelagic larval duration (Fig. B2 in Appendix B). For instance, the proportion of protected area where self- and external recruitment have equal contribution changed from 1% (10 km; Fig. B2a), to 17% (20 km; Fig. B2b), and then to 35% (40 km; Fig. B2c), and finally to over 50% (80 km; Fig. B2d) for species with pelagic larval duration of 8 days compare to 1% for 10 km, 1.5% for 20 km, 3.6% for 40 km, and then a 7.5% for 80 km for species with larval duration of 120 days.

Our results show that when spatial resolution is high within MPAs, highly nested subnetworks satisfy both conditions for network persistence: (1) to have a few self-persistent patches with higher local-retention, and (2) to be well connected to export and support the other patches.

### 4.3 Nested MPA networks: Interaction between MPA spacing and within-MPA connectivity

MPA spacing is a metric of interest as it can affect the amount of larval dispersal exchanged between MPAs and can alter the roles of external and self-recruitment. Our results revealed that the difference between the proportion of external and self-recruitment (*Ext – Self* in Fig. 4) response to MPA spacing varied with the proportion of protected area and with within-MPA spatial resolution. For instance, when the proportion of protected area is at 10%, the difference between the external and self-recruitment decreased as MPA spacing increased from 76 km to 250 km, but increased when spacing is further increased to 425 km for all levels of within-MPA spatial resolution and species pelagic larval durations values (Fig. 4a). Moreover, for the two high spatial resolution (10 km and 20 km), the difference between the external and self-recruitment is higher when MPA spacing is at its highest (425 km) compared to its lowest (76 km) value. However, the latter patterns change as the proportion of protected area increased. Surprisingly, for proportion of protected area of 20% (Fig. 4b) and 30% (Fig. 4c), the difference between external and self-recruitment monotonically decreased with MPA spacing for most within-MPA spatial resolutions. At proportion of protected area of 40%, the difference between external- and self-recruitment follows a response to MPA spacing similar to the case with 10% of protected area.

**Fig. 4.**
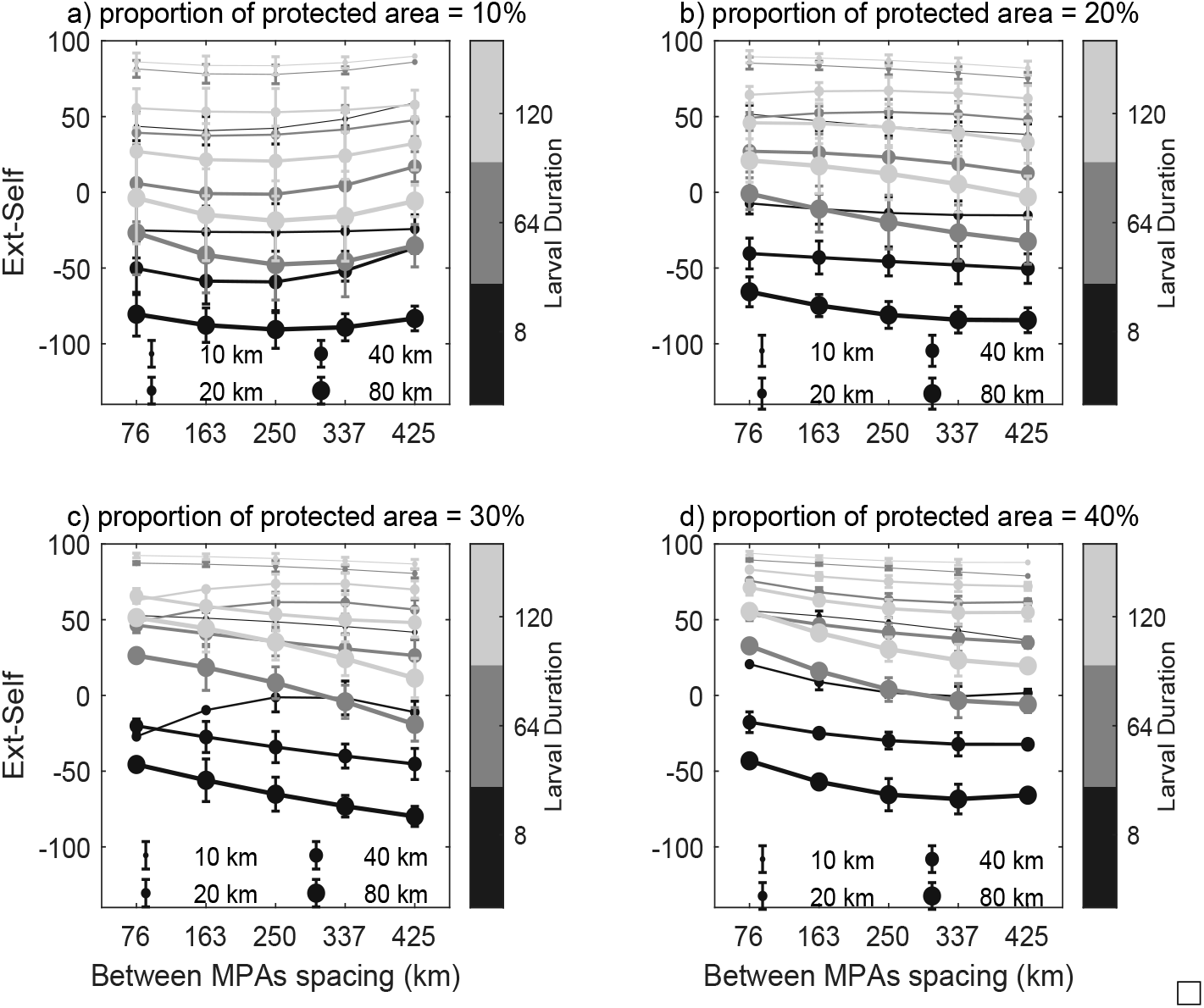
The difference between proportion of external- (*Ext*) and self-recruitment (*Self*) (*Ext – Self*) averaged among patches within-MPA and how it varies with MPA spacing, pelagic larval duration, and spatial resolution within MPAs for four proportions of protected area: (a) 10%, (b) 20%, (d) 30%, and (c) 40%. For each MPA spacing, the points and error bars represent the mean and standard deviation over MPA network replicates and spawning times (STs).

The minimal net reproductive rate (NRR) followed the inverse of response of difference between external and self-recruitment to MPA spacing, such that minimal NRR followed an increasing, decreasing, and a concave response when the difference between external and self-recruitment followed a decreasing, increasing, and convex response to MPA spacing, respectively. For pelagic larval duration of 8 days, the minimal NRR decreased with MPA spacing for high within-MPA spatial resolution when the proportion of protected area is 10% (10 km, Fig. 5a). However, as we increased the proportion of protected area the response changed from decreasing to increasing and then to decreasing (20% in Fig. 5b, 30% in Fig. 5c, and 40% in Fig. 5d). Intermediate (64 days) and longer (120 days) larval durations followed similar response to MPA spacing. However, as spatial resolution decreased, minimal NRR remained high across MPA spacing values for MPAs with spatial resolution of 20 km and showed a strong decreasing response for both 40 km and 80 km resolutions (Figs. 5c,d).

**Fig. 5.**
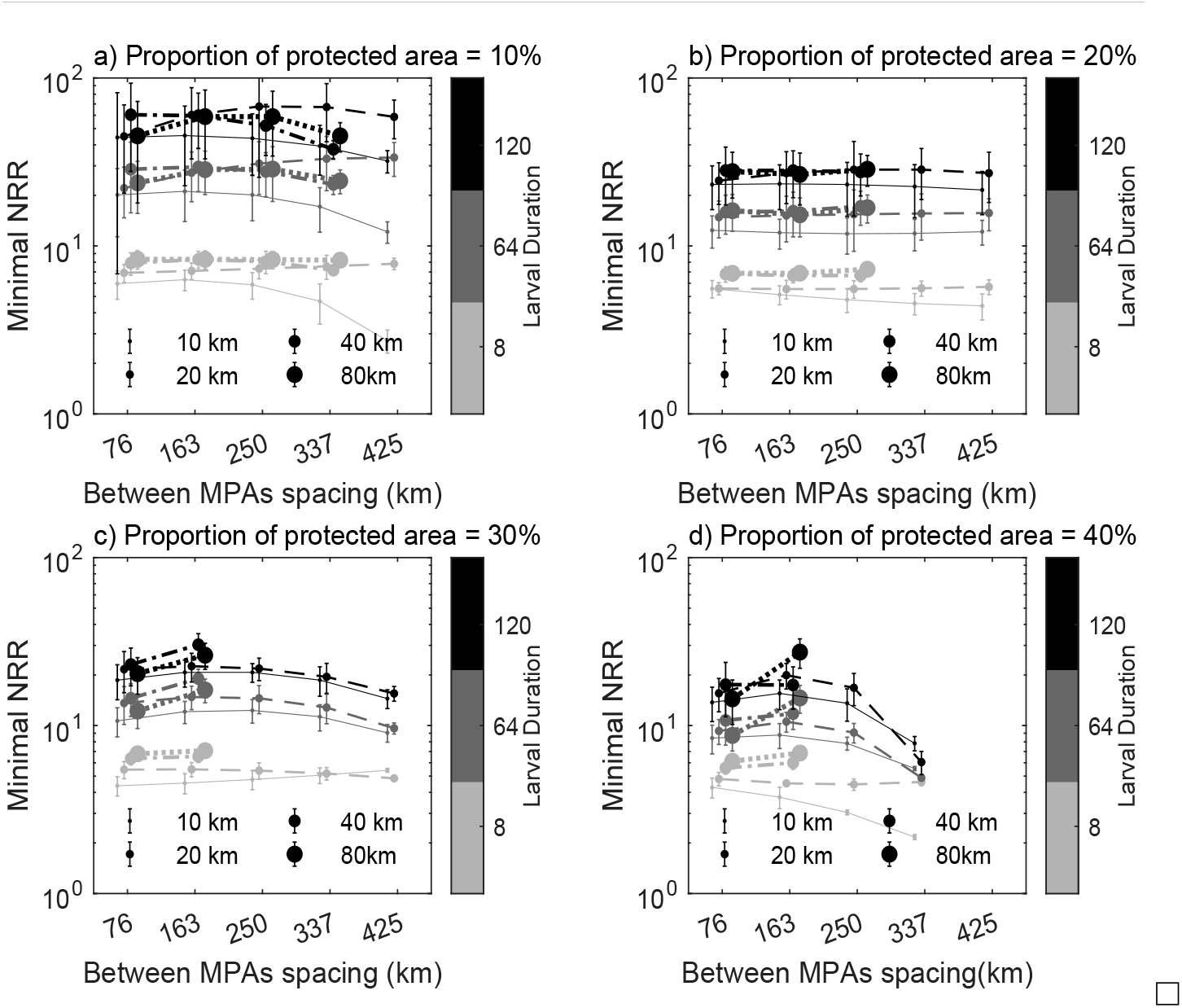
The minimal net reproductive rate (minimal NRR) and how it varies with MPA spacing for different pelagic larval durations and spatial resolution within MPAs (10 km, 20 km, 40 km, and 80 km) when proportion of protected area is: (a) 10%, (b) 20%, (c) 30%, and (d) 40%. For each MPA spacing value, dots and error bars represent the mean and standard deviation over MPA network replicates and spawning times (STs).

### 4.4 Metapopulation stability in MPA network with within-MPA connectivity

When we consider within-MPA connectivity as a nested subnetwork, each MPA contribution to stability 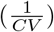 can be partitioned into its within-patch contribution to within- and between-MPA networks, both governed by dynamic connectivity within and between MPAs. Our results show how stability of overall MPA networks is affected by dynamic connectivity when within-MPA spatial resolution increased.

Within-MPA spatial resolution covaries with temporal variations in connectivity, and more specifically with its statistical moments including the mean, variance, covariance, and realized connections. Within-MPA spatial resolution decreased the mean and increased the variance of larval recruitment (Fig. 6), both changes resulted in reducing overall stability of MPA networks with increasing within-MPA spatial resolution when proportion of protected area is at 10% (Fig. 7). Moreover, since the mean and variance of connectivity decay exponentially with pelagic larval duration, we observed stronger response of short pelagic larval durations compared to longer pelagic larval duration (Fig. 6), and stability decreased minimally for longer compared to short pelagic larval duration (10% of protected area in Fig. 7).

**Fig. 6.**
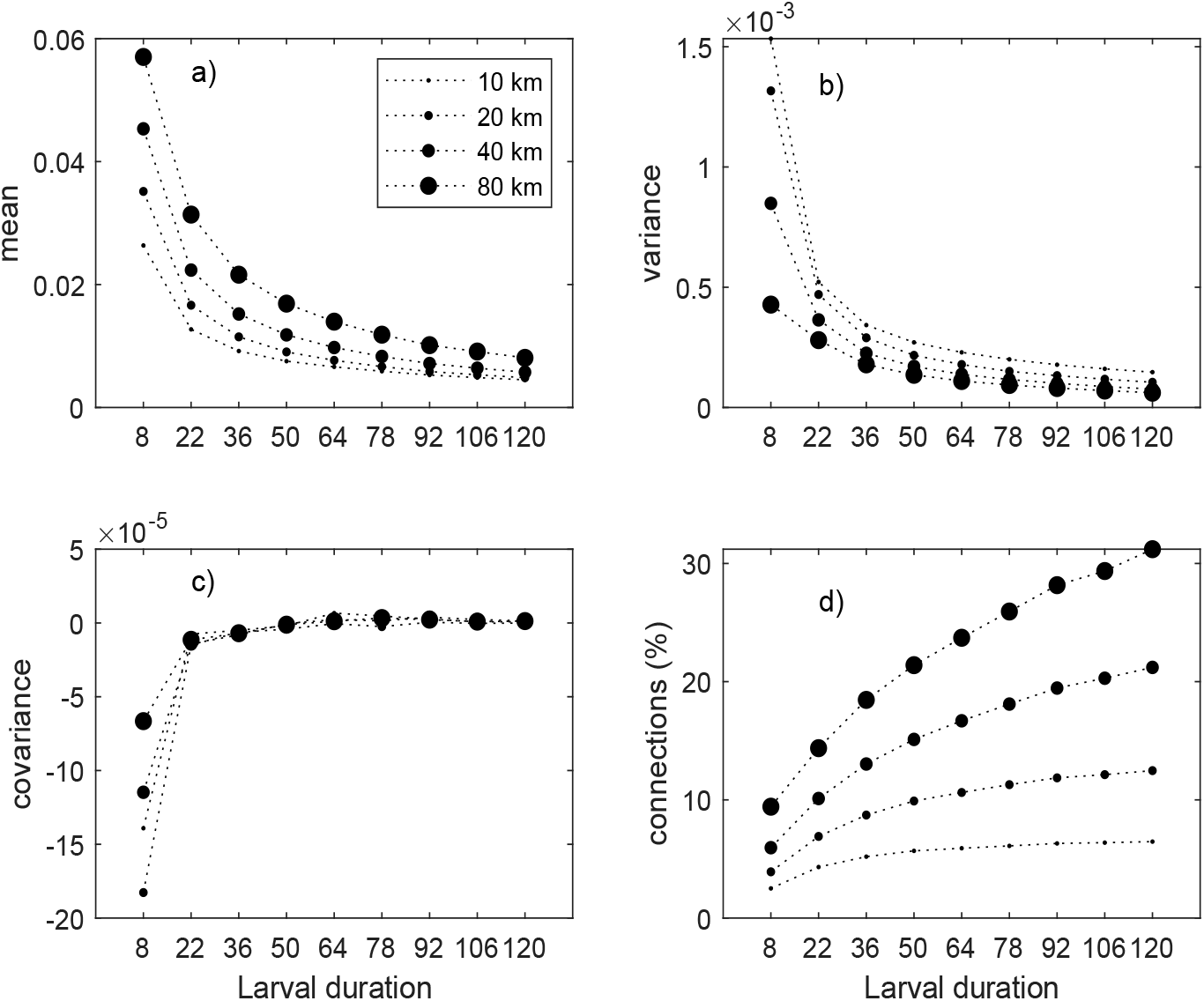
Response of statistical components of dynamic connectivity to pelagic larval duration and spatial spatial resolution within MPAs: (a) mean, (b) variance, (c) covariance, and (d) percentage of realized connections. For each pelagic larval duration, points represent the mean over MPA replicates and spawning times.

**Fig. 7.**
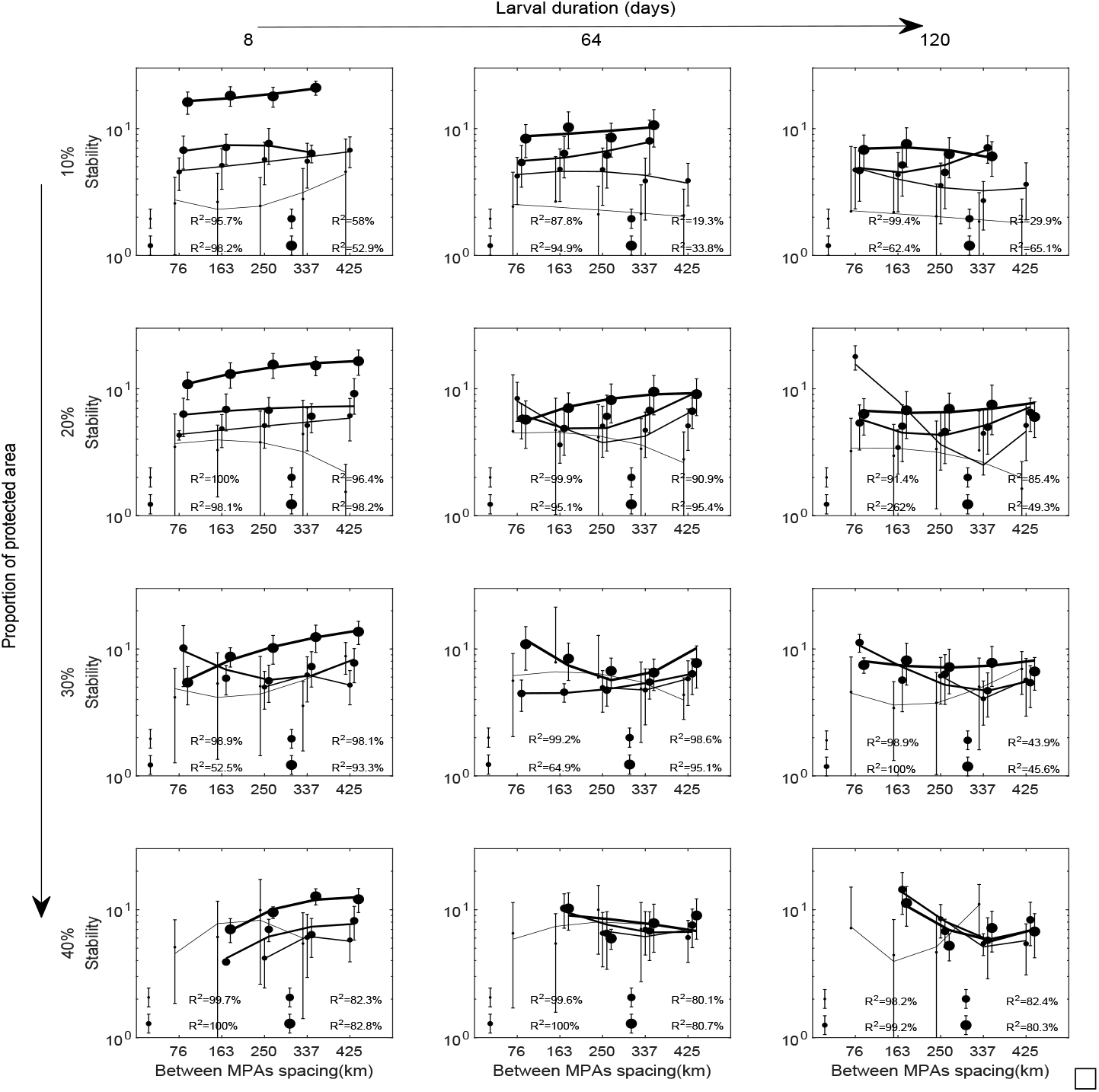
Mean MPA Network stability (1/*CV*) as a function of MPA spacing and within-MPA spatial resolution (dot size) for different pelagic larval durations (panel columns) and proportions of protected area (panel rows). Solid lines show best quadratic fit and error bars represent standard deviation over MPA replicates and spawning times

The response of metapopulation stability to MPA spacing within MPA networks varied with within-MPA spatial resolution, proportion of protected area, and pelagic larval duration. When the proportion of protected area is 10%, MPA spacing increased (decreased) network stability for short (long) pelagic larval durations regardless of the level of within-MPA spatial resolution. Increasing the proportion of protected area from 10% to 40%, increased (decreased) stability at shorter MPA spacing for high (low) spatial resolution especially with long pelagic larval durations.

The decreasing metapopulation stability in MPA networks with increasing within-MPA spatial resolution results from the combination of between- (*U*; Fig.2) and within- (*Ω*; Fig.2) MPA responses. These responses can be captured by the within-MPA population contributions to both within- 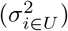 and between-MPA 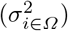 networks (eq. 7).

Metapopulation stability in MPA networks decreased with increasing MPA contribution 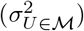, which itself increased with increasing difference between weighted variance contribution of populations to between- 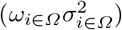 and within- 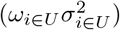 MPA networks. Larval recruitment when recruitment is from MPAs (*ω*_*i*∈*Ω*_) and within-MPA (*ω*_*i*∈*Ω*_) networks and variance contribution to between-MPA 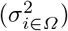 varied little compared to how variance contribution to within-MPA 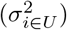 varied with within-MPA spatial resolution when the proportion of protected area is low (10% in Figs. B4 and B5). Within-MPA dynamics is what reduces MPA network stability: the higher the within-MPA spatial resolution, the lower the variance contribution of populations to within-compare to between-MPA networks (Fig. B4). The decrease of population contribution to the within-MPA variance translated into a stable within-MPA subnetworks and unstable between-MPA networks with increasing within-MPA spatial resolution.

## 5 Discussion

We studied the impact of connectivity within and between MPAs on metapopulation persistence in MPA network in relation to population growth from low density and to population stability at carrying capacity. Theories of MPA network design based on metapopulation dynamics typically focus on the study of population growth at low densities, mostly ignoring density-dependent processes (Burgess et al. 2014). However, the establishment of MPA networks not only aim to enable populations at low densities to recover, but also to promote the long-term stability of populations at higher abundances (Carr et al. 2017). When populations are near carrying capacity, their dynamics can be density-dependent (e.g. Hixon et al. 2012). Density-dependent dynamics involve mortality due intraspecific competition on both adults and new recruits, and both are affected by fluctuating larval dispersal/connectivity within and between MPAs. In general, our findings show connectivity dynamics within MPAs can promote the persistence of populations at both low and high densities. Specifically, our findings predict a lower minimum net reproductive effort (NRR) is required for growth of a populations at low densities and more stable dynamics of populations near carrying capacity.

In addition to the common assumption of low-density dynamics, most current MPA theories rely on temporally and spatially averaged dispersal (Burgess et al. 2014). Our work extends this current framework to account for spatially and temporally fluctuating connectivity interacting with density-dependent (meta)population growth across life stages. Differential responses to spatiotemporal dispersal of low and high density populations found in our study highlight the roles played by multiple statistical components of dynamic connectivity and their interaction with ecological processes such as intraspecific competition. At low densities, average larval recruitment is the most important predictor of population persistence. However, at high abundance, the temporal average, variance and covariance of larval recruitment interact to govern the stability of fluctuations around carrying capacity. This suggests that variance and covariance in recruitment can impact the long-term maintenance of (meta)populations in MPA networks. Recruitment is widely recognized as being spatially and temporally variable (Caffey 1985; Cowen and Sponaugle 2009) and our work suggests that the explicit consideration of this variability is important for understanding population dynamics.

Our study support results from other studies that incorporate spatiotemporal larval dispersal in MPA planning with the goal of designing networks that can be robust to environmental variations and climate change (e.g. Carr et al. 2017; Coleman et al. 2017). Similar to recent studies, we show the importance of pelagic larval traits, in particular pelagic larval duration (PLD), for predicting temporal patterns of dynamic connectivity (Bani et al. 2019). By explicitly considering larval dispersal within and between MPAs, we reveal how PLD interacts with the spatial structure and resolution of connectivity networks. For instance, larval recruitment for species with short PLD show strong spatial variation compared to species with long PLD (Fig. B1). This mapping between the pelagic traits and spatiotemporal properties of larval dispersal could predict the multiple and interacting effects of climate change on physical ocean currents and pelagic traits (Bani et al. in review).

MPAs can self-persist when local-retention is high enough to compensate for both larval post-settlement and adult mortality or persist as a network when at least one MPA is self-persistent and well connected to all other MPAs (Botsford et al. 2001; Armsworth 2002). Metapopulation persistence in MPA networks has commonly been formulated as a problem of MPA spacing and distance (Botsford et al. 2001; Kaplan et al. 2006; Moffitt et al. 2009). Studies have suggested that MPA size should be increased and spacing decreased in response to environmental variability, including variability that directly affect larval dispersal (Airamé et al. 2003; Shanks et al. 2003; Salm et al. 2006; Green et al. 2007; McLeod et al. 2009). Increasing size and decreasing spacing are both expected to increase the average larval recruitment regardless of environmental variability. However, these predictions are based on the assumption that larval recruitment is proportional to MPA size. This is clearly not the case in our results (Fig. 3c), which show that local retention in MPAs follows a bimodal - first decreasing then increasing - relationship with MPA size, especially for short larval duration (Fig. B1). It is more likely that increasing MPA size will decrease local retention for species with short PLD. The chances of recruiting back to a natal population decreases with MPA size within the MPA network, and MPA self-persistence thus decreases with MPA size.

We found that population growth and stability in MPA networks exposed to fluctuating connectivity are strongly dependent on within-MPA connectivity. Local retention under the assumption of well-mixed recruitment results from spatially-averaged self- and external larval recruitment within each MPA. By considering within-MPA dynamics, it is possible for an individual population within an MPA to persist when the spatially-averaged recruitment is not enough to compensate for post-settlement and adult mortality (Fig. 4b). In that context, the minimal NRR can reveal the interaction between within-MPA connectivity and MPA size for recovery when at low abundance: if all MPAs are self-persistent independently of distance, the minimal NRR should remain the same regardless of MPA distance. Otherwise, MPAs are dependent on external recruitment and within-MPA connectivity for network persistence. Despite the strong focus on connectivity in MPA theory, connectivity within MPAs has rarely been discussed. Yet, marine populations are commonly patchily distributed across relatively small spatial scales (Sale et al. 2006) and empirical evidence has shown that recruits into one population can come from nearby patches (Planes et al. 2009; Berumen et al. 2012). Our findings suggest that larval connectivity within MPAs can promote population persistence and stability.

Across our range of PLD values, we found that networks with a few large MPAs have greater self-recruitment than external recruitment, but networks with many small MPAs have greater external than self recruitment (Fig. 4d). Minimal NRR required for persistence in our smallest MPAs within networks set at a small proportion of protected area (¡10%) is smaller than in large MPAs. These same small MPAs are more self-persistent and export more larvae, thus subsidizing non-self-persistent MPAs and contributing to MPA network persistence. These results suggest that external recruitment from small MPAs allows reaching low minimum NRR with less overall proportion of protected area compared to large MPAs that rely on self-recruitment. When connectivity within MPAs is considered, small and adjacent sites within MPAs reduce minimal NRR variations among species with different PLD (Fig. 5b) compared to large MPAs with well-mixed dispersal that are sensitive to PLD (Fig. B2).

When population abundance is near carrying capacity, large MPAs are predicted to benefit the stability of MPAs networks when within-MPA connectivity is ignored. On the other hand, within-MPA aggregation of larval recruitment spatially decouples the dynamics of an individual MPA from the network, which destabilizes the MPA network. However, this increase in network-level variability resulting from within-MPA dynamics is alleviated by reducing both MPA size and spacing. For example, with 20% of the coastline protected, MPA network stability is robust to changes in within-MPA spatial resolution when MPAs are small and close to each other, but network stability becomes sensitive to within-MPA spatial resolution with increasing MPA spacing (Fig. 7).

MPA theory has long highlighted a perceived tradeoff between MPA spacing and size for achieving fisheries and conservation objectives (Hastings and Botsford 2003; Gaines et al. 2010): conservation objectives are predicted to be best achieved with large MPAs that are self-sustaining while fisheries objectives are best served by small MPAs that maximize spillover effects and larval export (e.g. Botsford et al. 2009). This tradeoff has led to the suggested implementation of MPA networks with moderately sized and spaced MPAs (Gaines et al. 2010). Our cross-scale analysis incorporating within-MPA connectivity predicts that many small and nearby MPAs can maintain high population abundance and greater network-level stability, and thus achieve conservation goals for sedentary species with dispersive larval stages, while simultaneously providing spillover effects and larval export to achieve fisheries goals. Our study suggests spatiotemporal fluctuations in connectivity can impact MPA network design through the consideration of within-MPA connectivity. The importance of such a cross-scale theory of MPA networks is likely to grow with ongoing climate change and associated increase in environmental variability.

## APPENDIX A Partitioning MPAs variance contribution into within-MPA patches variance contribution

A marine protected area contributes 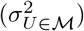 with the expression as follow:

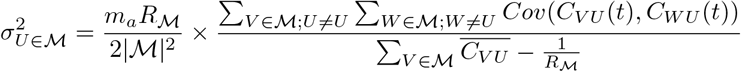

If we replace the connectivity between the MPAs (*C_VU_*(*t*), *C_WU_*(*t*), and 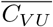) by their expression as a function of their within patches (Eq. 6):

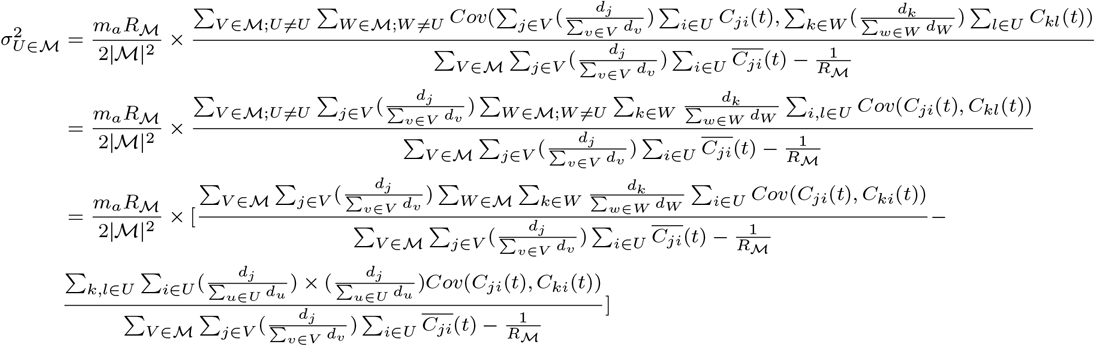

For simplicity and clarity, we assume that the patches do have similar size with equal net larval reproduction *d_i_* = *d_i_*, then 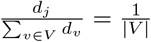:

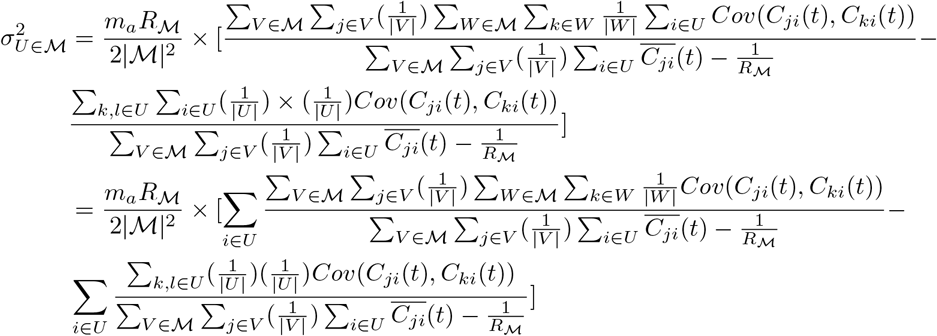

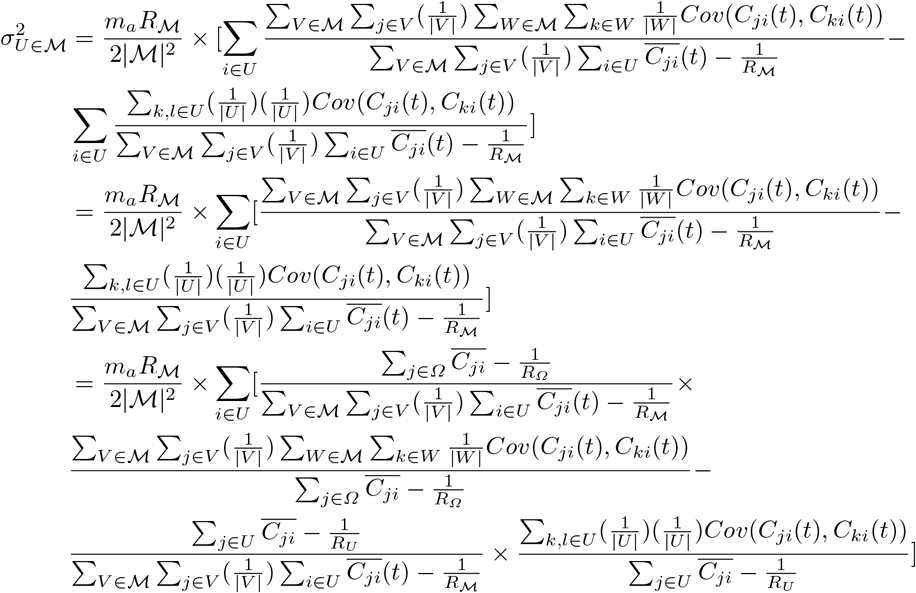

If we assume that MPAs nestedness is similar such that 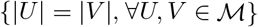, we can further simpify the expression into:

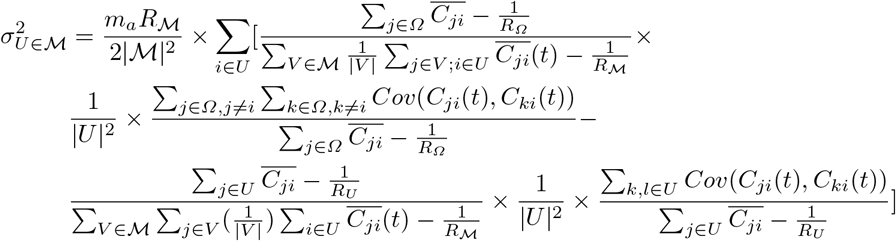

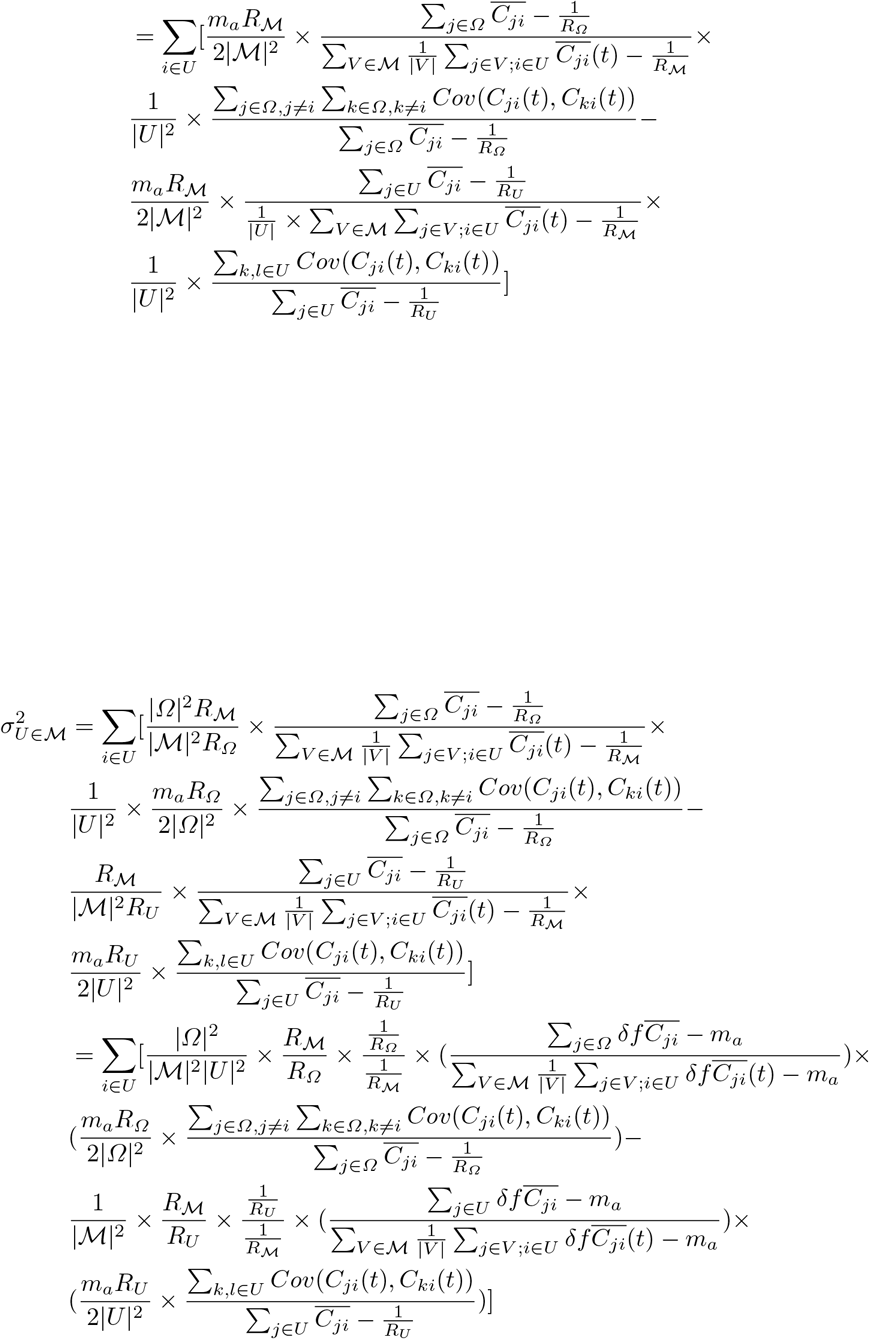

Then:

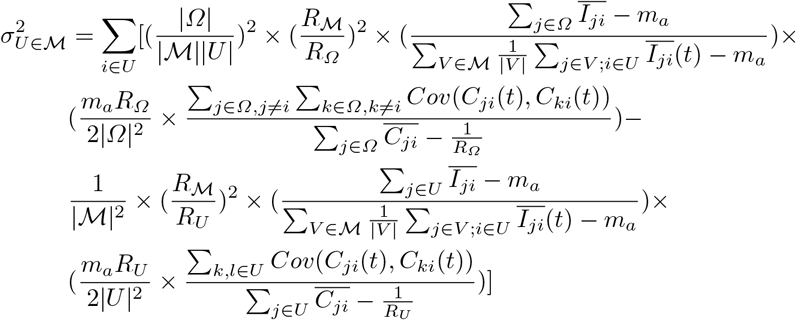

Therefore, if we let:

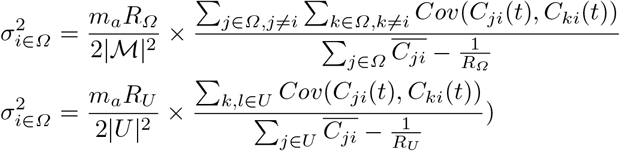

and:

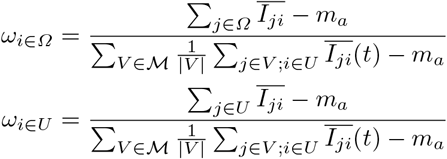

Thus, we can write the contribution in variance of MPA *U* to network stability as:

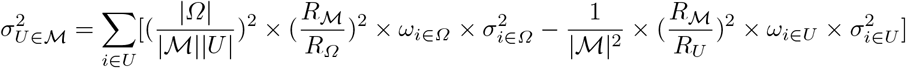

## APPENDIX B Supplementary Figures

**Fig. B1.**
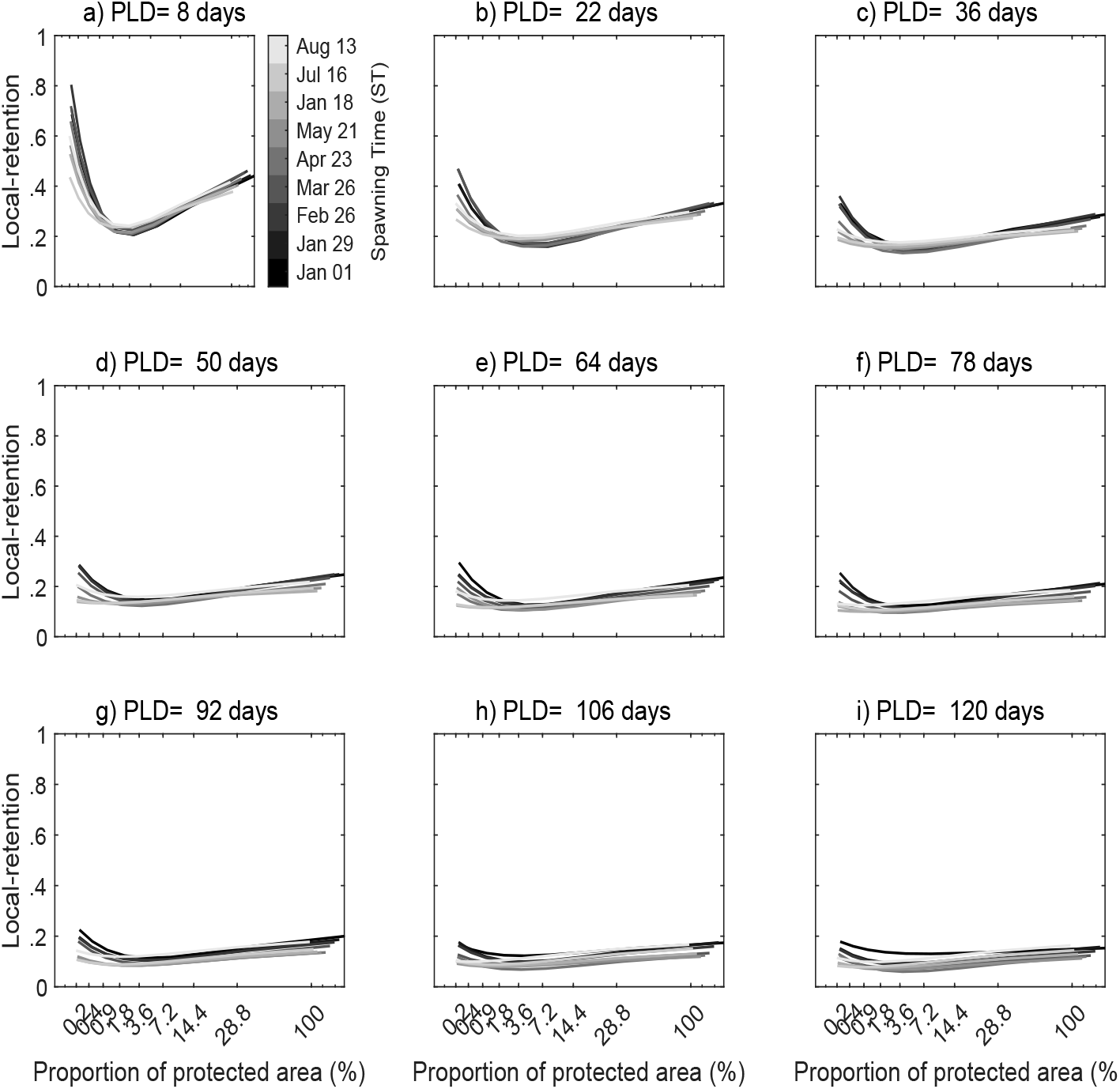
self-retention (probability of returning to natal population) of single MPAs and how it varies with the proportion of protected areas, pelagic larval duration (pld), and spawning time (colorbar).

**Fig. B2.**
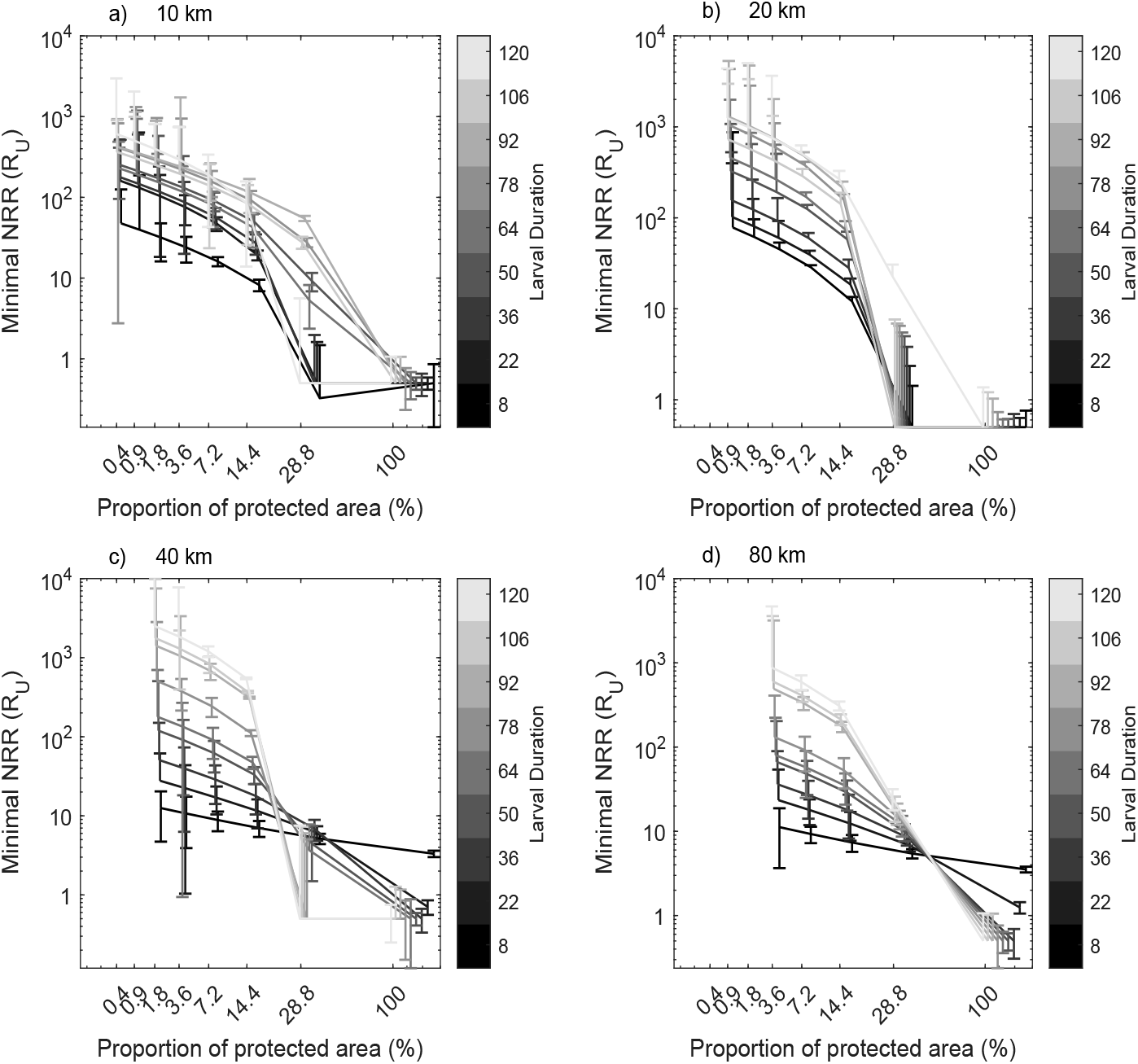
Minimal NRR required for persistence for the case of single MPA and how it varies with proportion of protected area and spatial resolution within-MPA: a) 10 km, b) 20 km, c) 40 km, and d) 80 km for different pelagic larval durations. For each proportion of protected area, the points and error bars represent averages and standard deviations for (b and c) different spawning times and MPA replicates. The spatial heterogeneity within-MPA reduced both minimal NRR required for persistence and differences between pelagic larval durations.

**Fig. B3.**
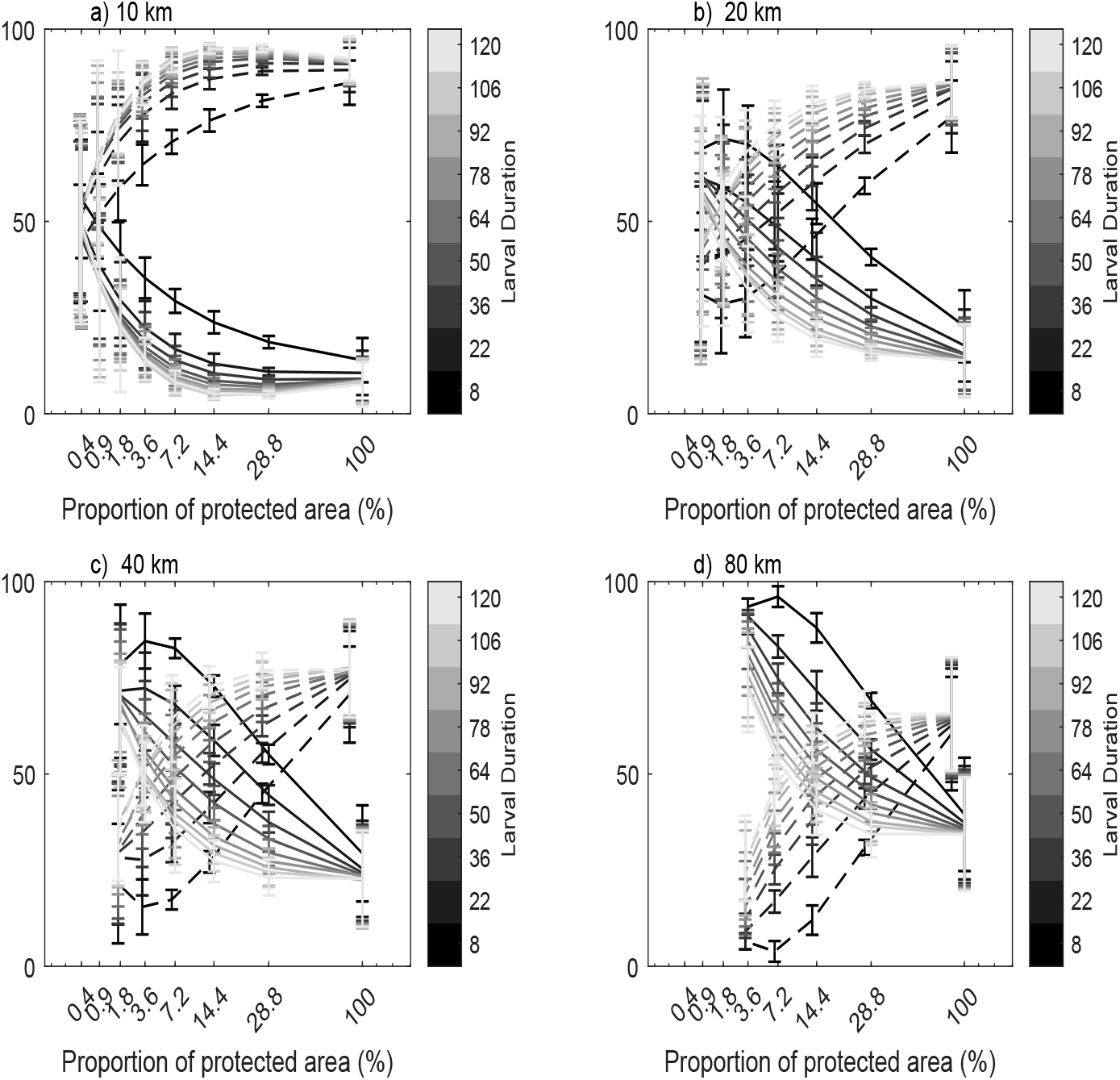
Self-recruitment (solid line) and external-recruitment (dashed line) of patches within-MPA in the case of single MPA and how it varies with proportion of protected area and spatial heterogeneity within-MPA: a) 10 km, b) 20 km, c) 40 km, and d) 80 km for different pelagic larval durations (color bar). For each proportion of protected area, the points and error bars correspond to fitted averages and actual standard deviations of different spawning times and MPA replicates, respectively. The data are fitted using quadratic functions.

**Fig. B4.**
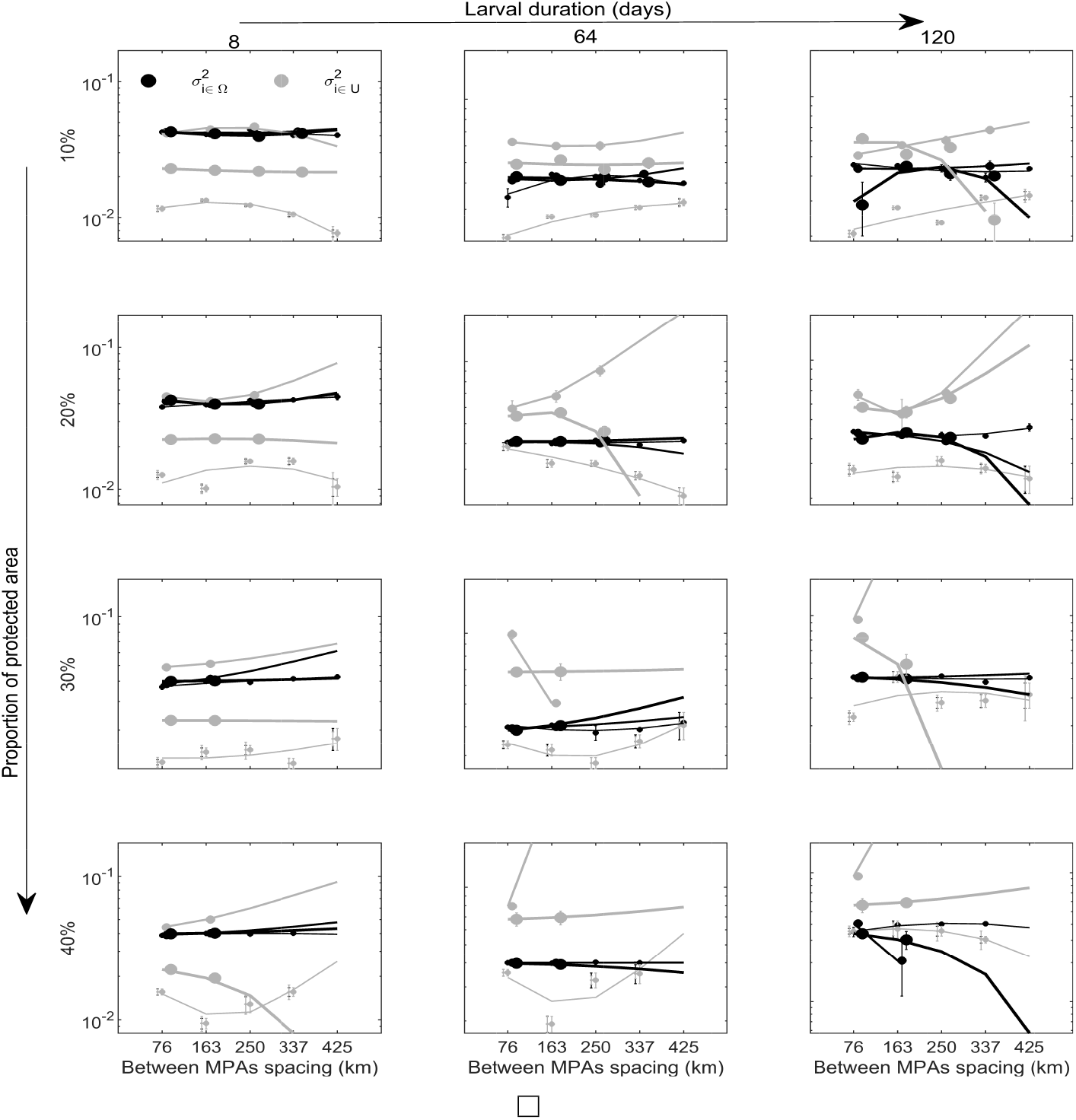
The variance contribution of patches within-MPA to whole 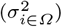 and within-MPA 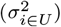 networks stability and how they vary with MPA spacing and spatial heterogeneity within-MPA (dots size: tiny 10 km, small 20 km, medium 40 km, and large 80 km) for different pelagic larval durations (columns: short 8 days, intermediate 64 days, and long 120 days) and proportion of protected areas (rows: 10%, 20%, 30%, and 40%). The lines are best quadratic fit, and points and error bars represent averages and standard deviations of different MPA replicates and spawning times. The within-MPA patch contribution to overall network 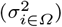 varied little with spatial heterogeneity within-MPA compared their variance contribution to within-MPA 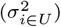 network stability regardless of MPA spacing especially at low proportion of protected areas (10%) and short pelagic larval duration.

**Fig. B5.**
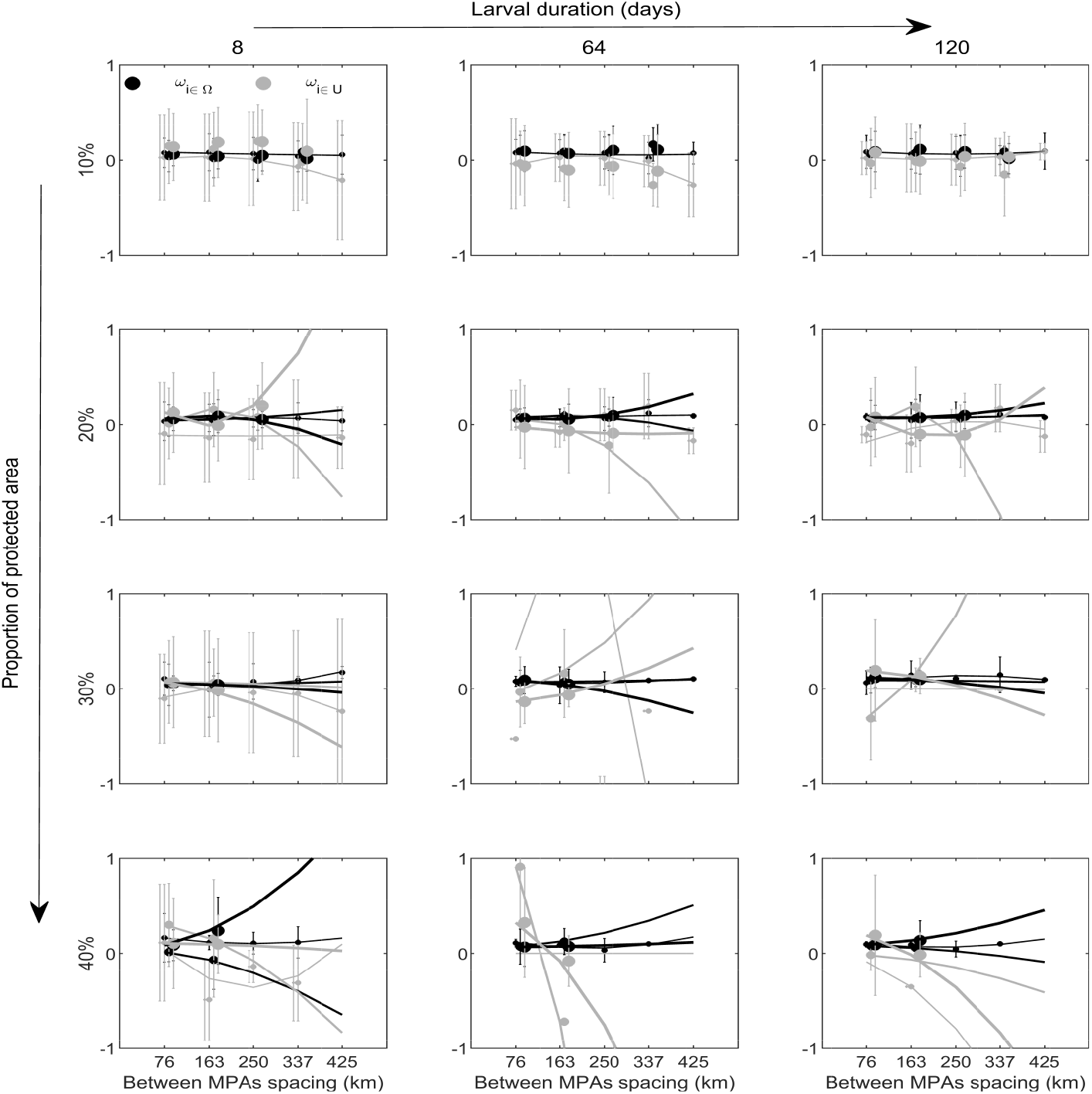
Relative larval recruitment when it is from whole (*ω*_*i*∈*Ω*_) and within-MPA (*ω*_*i*∈*u*_) networks (proportion in y-axis) and how they vary with MPA spacing (x-axis) and spatial heterogeneity within-MPA (dots size: tiny 10 km, small 20 km, medium 40 km, and large 80 km) for different pelagic larval durations (columns: short 8 days, intermediate 64 days, and long 120 days) and proportion of protected areas (rows: 10%, 20%, 30%, and 40%). The lines are best quadratic fit, and points and error bars represent averages and standard deviations of different MPAs replicates and spawning times, respectively. Larval recruitment from whole network varied little with spatial heterogeneity within-MPA compared larval recruitment from within-MPA network regardless of MPA spacing especially at low proportion of protected areas (10%) and short pelagic larval duration.

